# A vaccine built from potential immunogenic pieces derived from the SARS-CoV-2 spike glycoprotein

**DOI:** 10.1101/2020.09.24.312355

**Authors:** Jose Marchan

**Affiliations:** Experimental Medicine Centre, Venezuelan Institute for Scientific Research (IVIC). Apartado postal 20632, Caracas 1020-A, Venezuela., Email: /

**Keywords:** SARS-CoV-2, Spike, COVID-19, Immunoinformatics, Epitope, Vaccine

## Abstract

Coronavirus Disease 2019 (COVID-19) represents a new global threat demanding a multidisciplinary effort to fight its etiological agent—severe acute respiratory syndrome coronavirus 2 (SARS-CoV-2). In this regard, immunoinformatics may aid to predict prominent immunogenic regions from critical SARS-CoV-2 structural proteins, such as the spike (S) glycoprotein, for their use in prophylactic or therapeutic interventions against this rapidly emerging coronavirus. Accordingly, in this study, an integrated immunoinformatics approach was applied to identify cytotoxic T cell (CTC), T helper cell (THC), and Linear B cell (BC) epitopes from the S glycoprotein in an attempt to design a high-quality multi-epitope vaccine. The best CTC, THC, and BC epitopes showed high viral antigenicity, lack of allergenic or toxic residues, and suitable HLA-viral peptide interactions. Remarkably, SARS-CoV-2 receptor-binding domain (RBD) and its receptor-binding motif (RBM) harbour several potential epitopes. The structure prediction, refinement, and validation data indicate that the multi-epitope vaccine has an appropriate conformation and stability. Three conformational epitopes and an efficient binding between Toll-like receptor 4 (TLR4) and the vaccine model were observed. Importantly, the population coverage analysis showed that the multi-epitope vaccine could be used globally. Notably, computer-based simulations suggest that the vaccine model has a robust potential to evoke and maximize both immune effector responses and immunological memory to SARS-CoV-2. Further research is needed to accomplish with the mandatory international guidelines for human vaccine formulations.

**Graphical Abstract:** 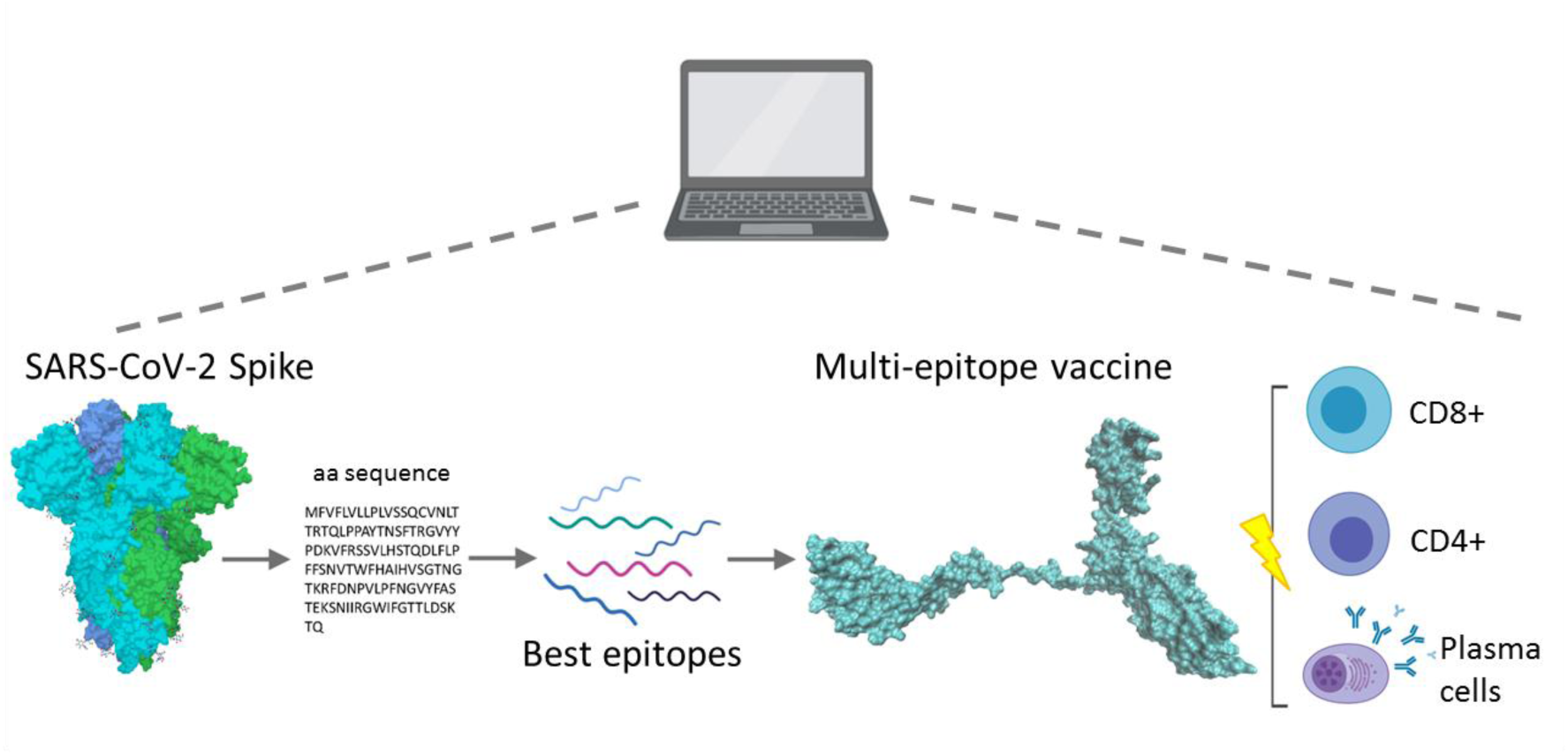

## 1. Introduction

On 31st December, 2019, a dramatic increase in the number of patients with a potentially fulminant respiratory disease was reported in Wuhan, China [1]. The etiological agent was eventually identified as a novel highly pathogenic betacoronavirus—severe acute respiratory syndrome coronavirus 2 (SARS-CoV-2) [2]. Shortly thereafter, SARS-CoV-2 caused an overwhelming wave of coronavirus disease 2019 (COVID-19) cases across Asia, Europe, Oceania, the Americas, and Africa, which led to the World Health Organization declared COVID-19 as a pandemic on 11th March, 2020 [1]. Unfortunately, at the time of this research, no clinically approved treatment is available to fight SARS-CoV-2, whose rapid spread has generated an explosive second wave of COVID-19 all over the world [1].

The SARS-CoV-2 outer membrane is decorated with several structural proteins, including the S glycoprotein, the membrane protein, and the envelope protein [3]. The S glycoprotein forms homotrimers containing both a receptor-binding domain (RBD) and a receptor-binding motif (RBM) [4]. The latter mediates contacts with human angiotensin-converting enzyme 2 (hACE2), thereby allowing SARS-CoV-2 entry into host cell [4]. This critical role in viral pathogenesis turns the SARS-CoV-2 S glycoprotein into an attractive target for vaccine development [5].

Multi-epitope vaccines designed from immunoinformatics tools could aid to elicit a protective immune response against SARS-CoV-2, as reported previously for other infectious agents [6,7]. In this regard, recent data indicate that the SARS-CoV-2 S glycoprotein harbours prominent immunologically active regions, which may serve as candidates for multi-epitope vaccine models [8-11]. Accordingly, the present study aimed to design a multiple-epitope vaccine construct against SARS-CoV-2 using for this purpose an integrated *in silico* approach.

## 2. Materials and methods

### 2.1. Protein sequence retrieval

Taking into account that the SARS-CoV-2 S glycoprotein represents the major target for vaccine development [5], the present work focused only on such viral spike. The complete amino acid sequence of the SARS-CoV-2 S glycoprotein was retrieved from Uniprot (http://www.uniprot.org/) in FASTA format (accession number: P0DTC2). Fig. 1 summarises the *in silico* experimental work.

**Fig. 1.**
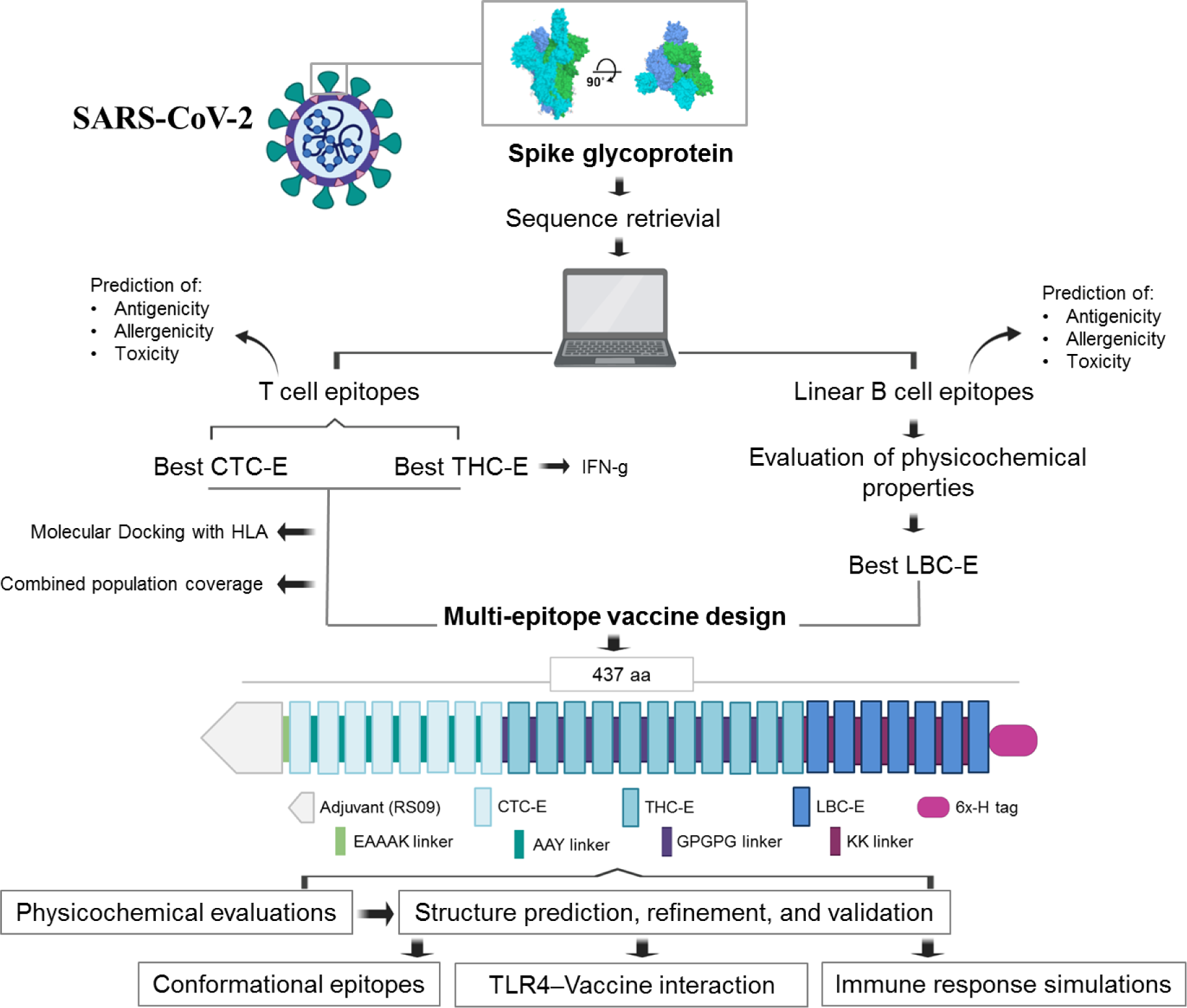
Overall experimental workflow (made with Biorender.com). Best epitopes predicted from the SARS-CoV-2 S glycoprotein were selected to design the multi-epitope vaccine construct, which was subjected to further in silico evaluations. CTC-E: cytotoxic T cell epitope. THC-E: T helper cell epitope. LBC-E: linear B cell epitopes. IFN-g: Interferon gamma. aa: amino acid. 6x-H: polyhistidine tag.

### 2.2. Prediction of allergenicity, toxicity, and viral antigenicity

Prior to clinical trials, safety evaluations of new vaccine candidates must include allergenicity and toxicity studies to avoid possible harm to humans due to the vaccine active ingredients [12]. Therefore, potential epitopes from the S glycoprotein were subjected to allergenic evaluation using two webservers: AlergenFP (http://ddg-pharmfac.net/AllergenFP/index.html) [13], and Algepred (www.imtech.res.in/raghava/algpred/) [14], whereas toxicity was predicted using the ToxinPred server (http://crdd.osdd.net/raghava/toxinpred/) [15]. Finally, viral antigenicity was calculated from the Vaxijen server (threshold: 0.5) (www.ddg-pharmfac.net/vaxijen/) [16].

### 2.3. Immunogenicity

Vaccine-mediated protection is intrinsically related to the activation of the cell-mediated and humoral responses [17]. The cell-mediated response is mainly based on the clonal expansion of human leukocyte antigen (HLA) class I-restricted CD8+ cytotoxic T cells (CTC), which are critical to antiviral defence during acute infection, and HLA class II-restricted CD4+ T helper cells (THC), which are necessary for the generation of specific antiviral antibodies when they crosstalk with B cells (BC) [18]. The humoral response, in contrast, involves the production of antibodies by BC [18]. Therefore, CTC, THC, and BC epitopes were predicted and the best were selected for the final vaccine design (Fig. 1). To achieve this aim, multiple prediction tools were used to improve the rate of true positives.

#### 2.3.1. Prediction of CTC and THC epitopes

Peptides that interact with HLA class I (HLA-I) and HLA class II (HLA-II) molecules commonly have 9 and 15 amino acids in length, respectively [18]. In consequence, 9-mer and 15-mer peptides were considered in this work as CTC and THC epitopes, respectively (Fig. 1). These epitopes were identified using the Immune Epitope Database and Analysis Resource (IEDB-AR) (http://tools.immuneepitope.org/main/) [19].

To cross-validate binding peptides to HLA molecules, several methods were applied. In this regard, the 9-mer binding peptides to HLA-I were predicted using the artificial neural network (ANN) method [20], the Consensus method [21], and the NetMHCpan method [22]. The prediction of the 15-mer binding peptides to HLA-II was performed using the Consensus method [23], the NetMHCIIpan method [24], and the SMM-align method [25]. Peptides were selected according to their percentile rank—peptides with a small percentile rank have high affinity by HLA alleles [21]. This percentile rank is generated on IEDB-AR by comparing the IC50 of each predicted peptide against random peptides from SWISSPROT database.

In addition, binding peptides to HLA-II were also chosen by their potential to induce interferon-gamma (IFN-g) (Fig. 1), which was evaluated using the IFNepitope server (http://crdd.osdd.net/raghava/ifnepitope/) [26].

#### 2.3.2. Prediction of linear BC epitopes

BCPRED (http://ailab.ist.psu.edu/bcpred/) [27] was used to predict linear BC epitopes based on several physicochemical properties: hydrophilicity, flexibility, accessibility, and antigenicity propensity (threshold = 1 for each parameter). Simultaneously, the S glycoprotein amino acid sequence was also subjected to iBCE-EL (http://thegleelab.org/iBCE-EL/) [28] and BepiPred-2.0 (http://www.cbs.dtu.dk/services/BepiPred/) [29] for additional predictions of linear BC epitopes.

#### 2.3.3. Three-dimensional (3D) interaction between HLA alleles and viral peptides: Molecular docking

To evaluate the presentation of the best epitopes in the context of HLA molecules, a molecular docking study was conducted (Fig. 1). Taking into account that HLA-C*06:02 and HLA-DRB1*01:01 were predicted as common interacting HLA alleles, they were selected for this purpose.

The molecular docking simulation process is summarised as follows. First, the best 3D structure of each epitope (9-mer and 15-mer peptides) was obtained from PEPFOLD server (https://bioserv.rpbs.univ-paris-diderot.fr/services/PEP-FOLD3/) [30]. Second, the 3D structure of HLA-C*06:02 and HLA-DRB1*01:01 were downloaded from Protein Data Bank (PDB) (accession numbers: 5W6A and 1AQD, respectively). Third, the receptor-ligand docking simulation was performed using the PatchDock server (http://bioinfo3d.cs.tau.ac.il/PatchDock/), which is based on shape complementarity principles [31]. Fourth, the receptor-ligand complexes were visualized with the VMD software (Version 1.9.3) [32]. Finally, the best allele-epitope complexes were selected based upon visual inspection and the PatchDock criteria, as well as they were compared with their respective control peptides (co-crystal ligands of HLA-C*06:02 and HLA-DRB1*01:01).

### 2.4. Design of the multi-epitope vaccine against SARS-CoV-2

High potential CTC, THC, and linear BC epitopes were selected to generate the amino acid sequence of the multi-epitope vaccine. The CTC and THC epitopes were linked together using AAY and GPGPG linkers, respectively, whereas linear BC epitopes were connected by KK linkers (Fig. 1). Moreover, a TLR4 agonist, known as RS09 (Sequence: APPHALS) [33], was added as an adjuvant at the N-terminus by using an EAAAK linker (Fig. 1). This adjuvant was chosen because of two relevant points: 1) the coronaviruses S glycoprotein may bind and activate TLR4 [34]; 2) a previous report has shown that RS09 favours a good binding affinity between multi-epitope vaccines and TLR4 [35]. For future purification and identification purposes, a polyhistidine-tag (6x-H tag) was included at the C-terminus (Fig. 1).

#### 2.4.1. Physicochemical evaluation and general studies

The ProtParam tool (https://web.expasy.org/protparam/) [36] was used to examine relevant physiochemical parameters of the multi-epitope vaccine. To reconfirm its viral antigenicity and lack of allergenicity and toxicity, the web tools described in section 2.2 were applied. In addition, the vaccine solubility was predicted using the SOLpro server (http://scratch.proteomics.ics.uci.edu/) [37].

#### 2.4.2. Structure prediction, refinement, and validation

PSIPRED (http://bioinf.cs.ucl.ac.uk/psipred/) [38,39] and GalaxyWEB (http://galaxy.seoklab.org/) [40,41] were utilized to predict the secondary and tertiary structure, respectively, of the multi-epitope vaccine construct. The best model was refined with GalaxyWEB [40,41] whereas its visualization was obtained from the VMD software (Version 1.9.3) [32]. The data validation was performed using PROCHECK (https://servicesn.mbi.ucla.edu/PROCHECK/) [42] and ProSA-web (https://prosa.services.came.sbg.ac.at/prosa.php) [43].

#### 2.4.3. Prediction of conformational BC

Conformational epitopes of the multi-epitope vaccine construct were predicted from Ellipro (http://tools.iedb.org/ellipro/) [44], which represents the protein structure as an ellipsoid and calculates protrusion indexes for protein residues outside of such ellipsoid [44]. Minimum levels of 0.8 and a distance of 6.0 Å were applied. The epitopes were visualized with the VMD software (Version 1.9.3) [32] to illustrate their position and 3D structure.

#### 2.4.4. Innate immune receptor-Vaccine interaction: Molecular docking

Since TLR4 may serve as a sensor for the recognition of coronaviruses S glycoproteins [34], this germline-encoded pattern recognition receptor was selected for the docking study. The 3D structure of TLR4 was obtained from PDB (accession number: 4G8A). The refined model of the multi-epitope vaccine was used as a ligand. The TLR4-Vaccine docking simulation and its 3D visualization were performed with Patchdock [31] and VDM [32], respectively.

#### 2.4.5. In silico immune response simulations

To further characterize the potential immune response of the multi-epitope vaccine, immune simulations were performed using the C-ImmSim server (http://150.146.2.1/C-IMMSIM/index.php) [45]. Three injections were applied four weeks apart as described previously [46]. Furthermore, 12 injections were applied four weeks apart to simulate repeated exposure to potential immunogen. The Simpson index D was used to interpret the diversity of the immune response.

#### 2.4.6. Population coverage

Global population coverage of the multi-epitope vaccine construct was calculated from IEDB-AR (http://tools.iedb.org/population/) [47,19]. The HLA allele genotypic frequencies available on IEDB-AR were obtained from Allele Frequency Database (AFD) (http://www.allelefrequencies.net/). At present, AFD contains allele frequencies from 115 countries and 21 different ethnicities (http://www.allelefrequencies.net/). Those 115 countries were selected and the HLA-I and HLA-I interacting alleles predicted in this work were included to perform an HLA combined analysis. The results were shown on a world map using Rstudio software (Version 3.5.3) [48].

## 3. Results

### 3.1. Prediction of T cell epitopes

A total of 47 T cell epitopes were predicted; however, 8 CTC and 11 THC epitopes were identified as the best (Table 1). These 19 epitopes showed a potent viral antigenicity— ranging from 0.63 to 1.52—and lack of allergenic or toxic residues in their sequences (Table 1). Moreover, THC epitopes were characterized by their potential capability to induce IFN-g (Table 1). Although “EGFNCYFPLQSYGFQ” (E47 in Table 1) could be categorized as a strong potential THC epitope, it was identified as a probable inductor of toxicity. Therefore, this epitope was not included in the amino acid sequence of the multi-epitope vaccine.

**Table 1.**
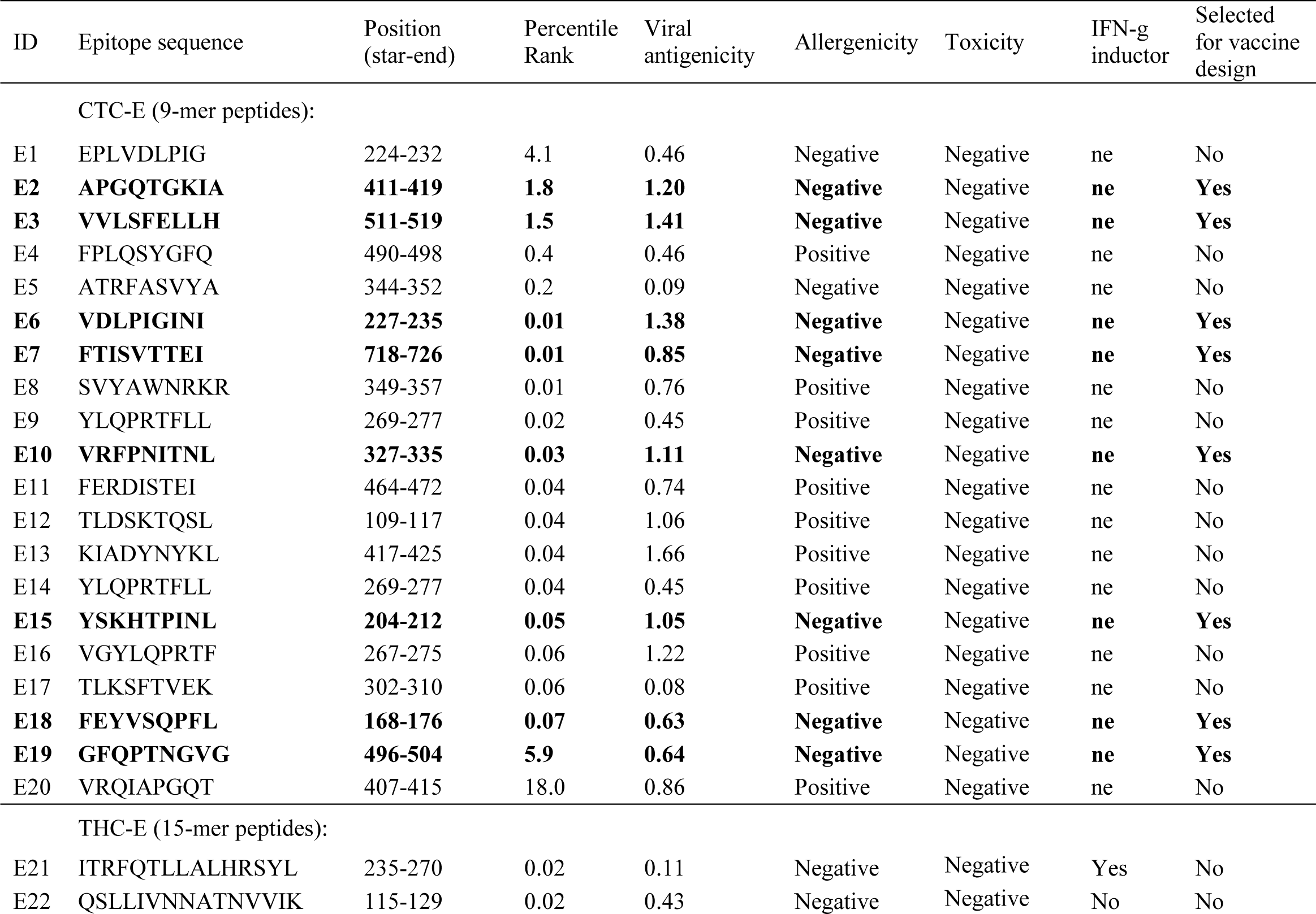
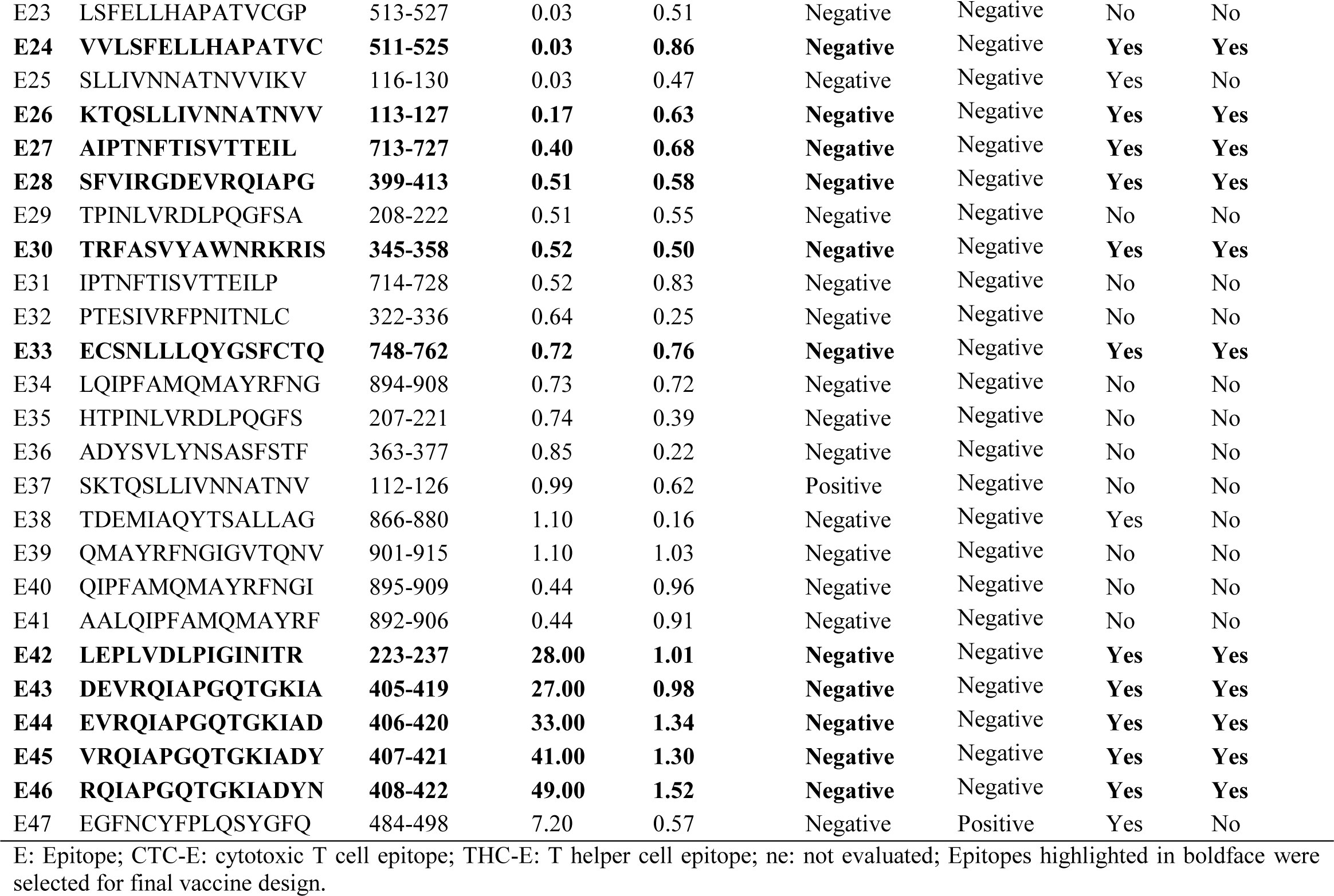
Evaluation of potential T cell epitopes.

#### 3.1.1. HLA-I and HLA-II interacting alleles

The selected CTC epitopes (Table 1) showed promiscuous affinity by several HLA-I alleles, including A*01:01, A*02:01, A*03:01, A*11:01, A*23:01, A*25:01, A*30:01, A*68:01, A*74:01, B*07:02, B*08:01, B*13:01, B*13:02, B*14:02, B*15:01, B*15:02, B*18:01, B*27:02, B*35:03, B*40:01, B*58:01, C*01:02, C*02:02, C*02:09, C*03:02, C*03:03, C*03:04, C*04:01, C*05:01, C*06:02, C*07:01, C*08:01, C*12:02, C*12:03, C*14:02, C*15:02, C*16:01, and C*17:01. Likewise, the selected THC epitopes (Table 1) showed common interaction with the following HLA-II alleles: DRB1*01:01, DRB1*01:03, DRB1*07:01, DRB1*15:01, DRB1*01:20, DRB3*02:02, DRB4*01:01, and DRB5*01:01.

### 3.2. Prediction of linear BC epitopes

A total of 10 linear BC epitopes of varying amino acid lengths were predicted (Table 2). Most of the epitopes showed robust viral antigenicity (≥0.5), as well as, they were identified as non-allergenic and non-toxic (Table 2). However, only 7 epitopes were selected for the vaccine design due to they were predicted simultaneously by 3 different web tools (BCPRED, iBCE-EL, and BepiPred-2.0) (Table 2). Interestingly, overlapping residues were observed between some linear BC and T cell epitopes.

**Table 2.**
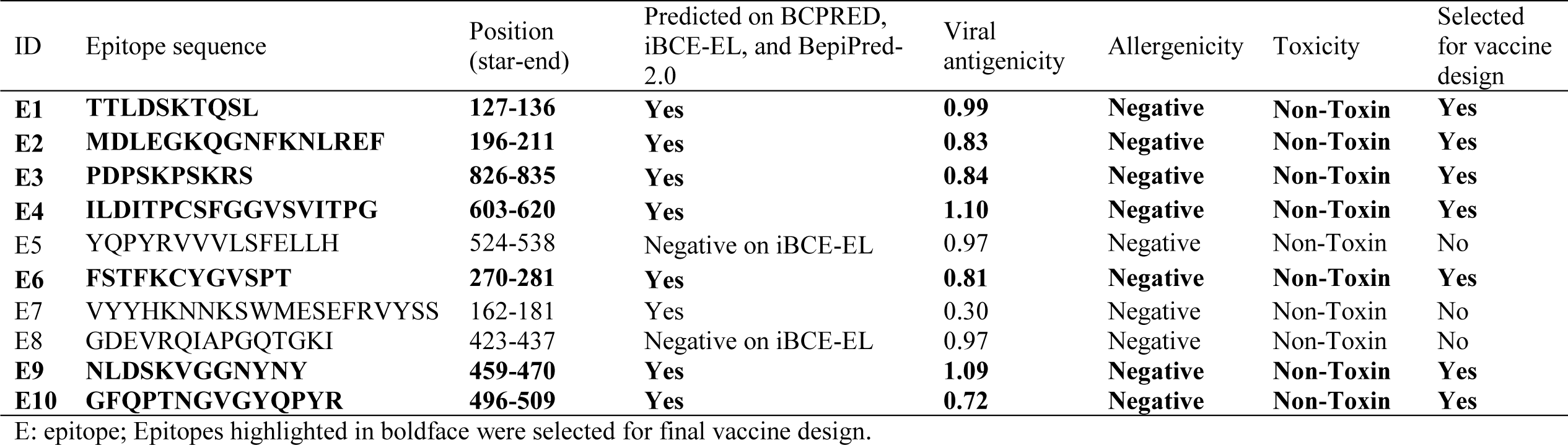
Evaluation of potential linear B cell epitopes.

### 3.3. HLA allele-viral peptide interaction: Molecular docking

To evaluate the presentation of the best epitopes in the context of HLA, molecular docking simulations were conducted. For this purpose, HLA-C*06:02 and HLA-DRB1*01:01 were chosen as representative alleles.

HLA-I and HLA-II alleles were docked with CTC and THC epitopes, respectively, using the Patchdock server, which has been recently applied to successfully dock epitopes from SARS-CoV-2 into immune cell receptors [10]. The HLA allele-viral peptide complexes showed high geometric shape complementarity scores (>6000) [52] similar to controls (Table 3). The inspection on VDM software allowed observing different binding patterns wherein viral peptides rightly interact with the active site residues of the HLA groove in a similar way to control peptides (Fig. 2). Moreover, several viral peptides (e.g., E15 and E33) formed a bulge that projected from their respective HLA allele (Fig. 2B and 2C), which may suggest a more direct interaction with the T cell receptor [18].

**Table 3.**
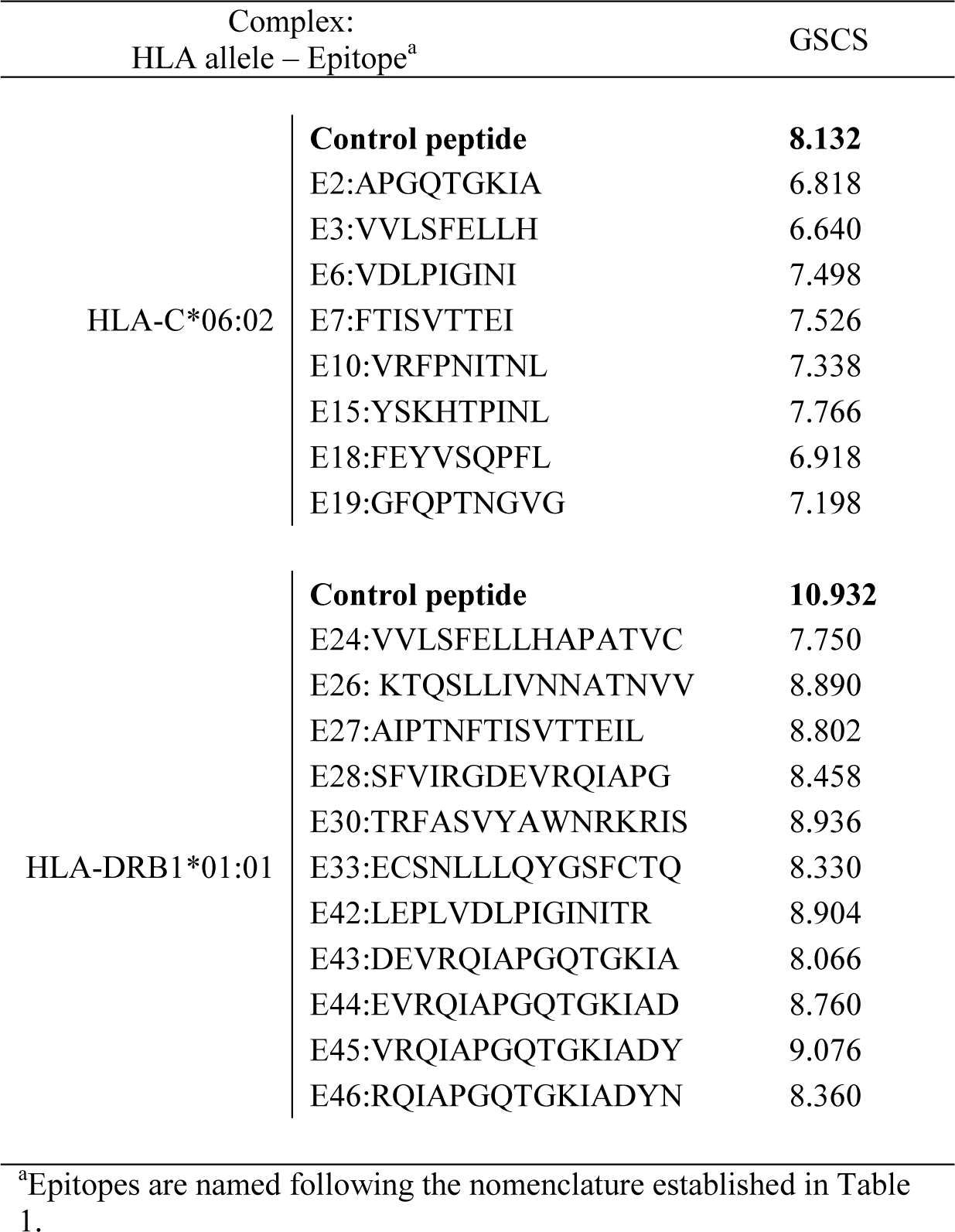
Patchdock score based on geometric shape complementarity scores (GSCS) of the HLA allele-epitope complexes.

**Fig. 2.**
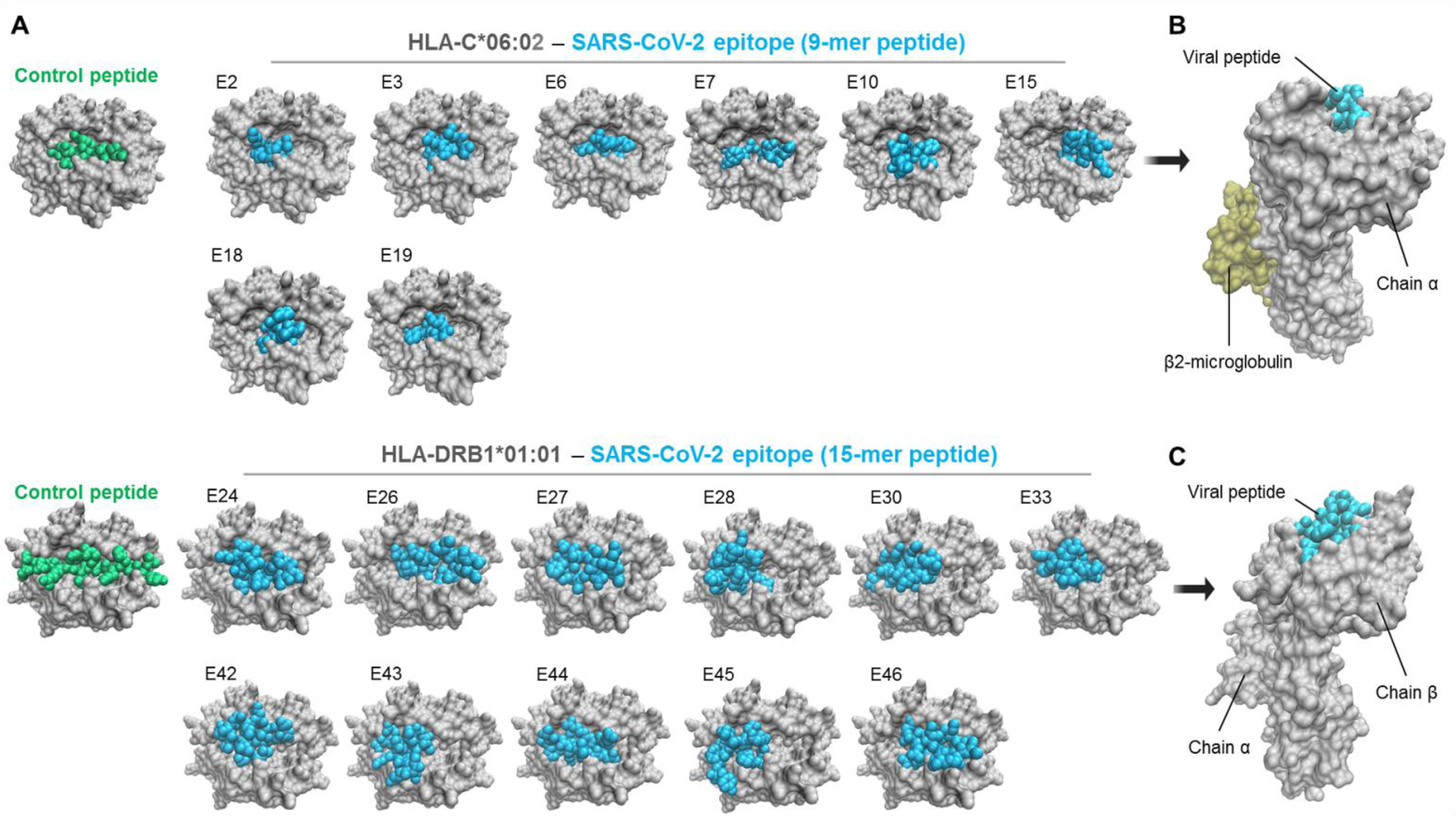
Screenshots of the HLA-viral peptide complexes. (A) Top view of HLA-C*06:02 and HLA-DRB1*01:01 presenting 9-mer and 15-mer viral peptides, respectively. (B-C) Representative lateral views of HLA alleles interacting with viral peptides. Of note, peptides formed a bulge that project from the HLA groove. Epitopes are named according to the nomenclature established in Table 1. HLA alleles, viral peptides, and control peptides are shown in grey, cyan, and green, respectively.

### 3.4. Design of the multi-epitope vaccine against SARS-CoV-2

#### 3.4.1. General evaluations

To design the amino acid sequence of the multi-epitope vaccine, a total of 26 epitopes (8 CTC, 11 THC, and 7 LBC epitopes) were organized using several linkers (Fig. 1). This sequence is constituted by 437 amino acid residues (Fig. 1).

Of particular note, 6 epitopes selected for the vaccine design (E2, E19, E43, E44, and E45 in Table 1; E10 in Table 2) harbour residues that are usually involved in the interaction between the SARS-CoV-2 S glycoprotein and hACE2 [49-51]. For instance, N501—which is present in the amino acid sequence of E19 (Table 1) and E10 (Table 2)—has been recently described as one of the critical hACE2-binding residues in SARS-CoV-2 [51].

The vaccine showed a strong viral antigenicity (0.64), as well as neither allergenic nor toxic residues were observed in its amino acid sequence. Furthermore, the physicochemical properties examined with the ProtParam tool, including molecular weight, theoretical pI, amino acid composition, atomic composition, extinction coefficient, estimated half-life, instability index, aliphatic index, and GRAVY, were computed as conventional results (Table 4).

**Table 4.** Physicochemical parameters of the multiple-epitope vaccine construct.

| Parameter | Value | Comment |
| --- | --- | --- |
| Number of aa | 437 | Suitable |
| Molecular Weight | 46kDa | Average |
| Theoretical pI | 9.61 | Slightly basic |
| Negatively charged residues (N + Q) | 25 |  |
| Positively charged residues (R + K) | 44 | Charged positive |
| Extinction coefficient (at 280 nm in water) | 32570 M<sup>-1</sup>cm<sup>-1</sup> | - |
| The instability index (II) | 26.02 | Stable |
| Aliphatic index | 74.58 | Thermostable |
| GRAVY | -0.298 | Hydrophilic |
| Solubility | 0.703269 | Soluble |
| Estimated half-life: |  | Satisfactory |
| Mammalian reticulocytes (in vitro) | 4.4 hours |  |
| Yeast (in vivo) | >20 hours |  |
| *Escherichia coli* (in vivo) | >10 hours |  |

aa: amino acid; GRAVY: Grand average of hydropathicity

#### 3.4.2. Structure prediction, refinement, and validation

The 437 amino acid long vaccine construct was analysed using the PSIPRED server to predict its secondary structure, which identified 316, 70, and 51 amino acids forming coil, helix, and strand regions, respectively (Fig. 3A).

**Fig. 3.**
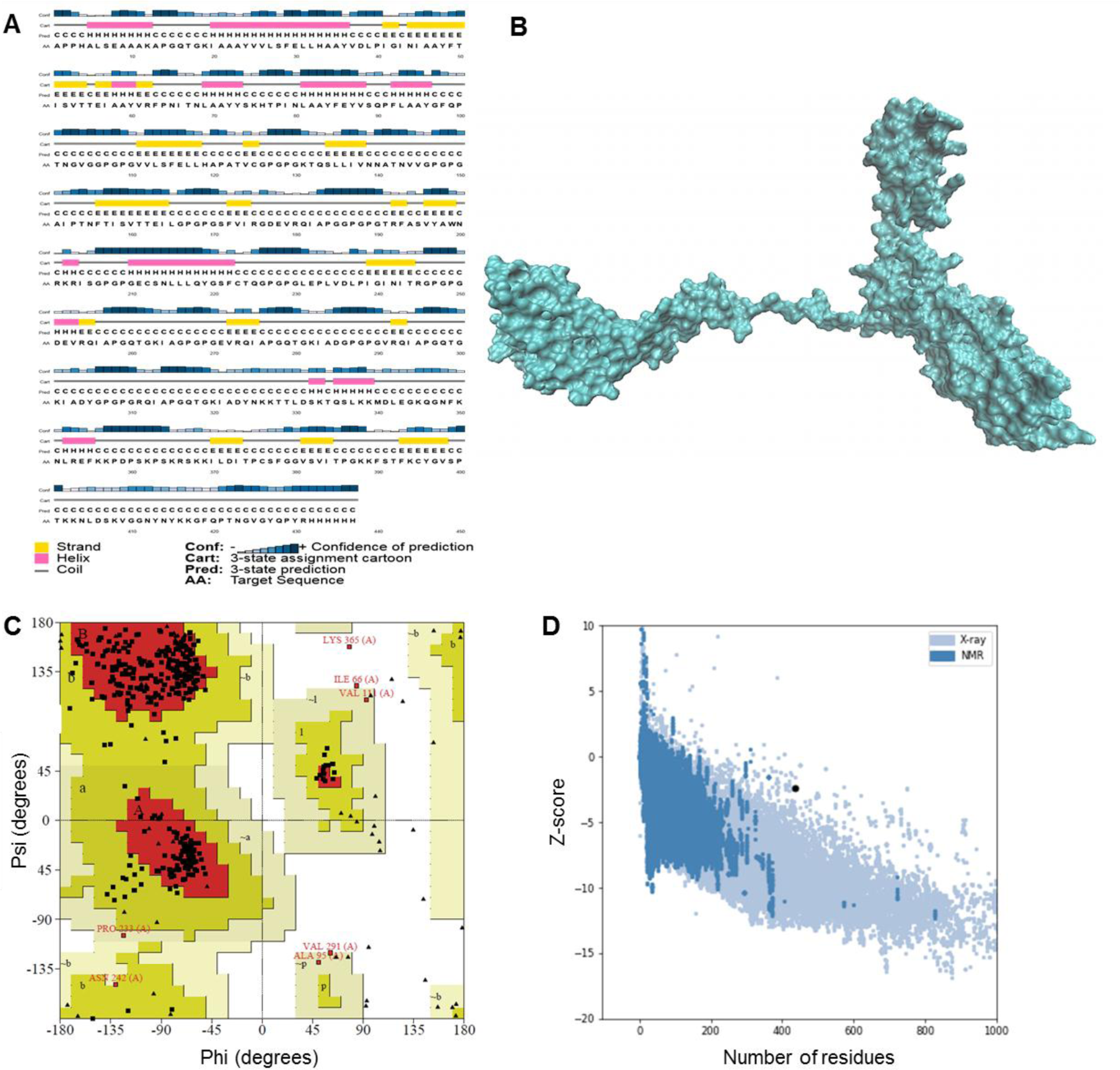
Structure predictions and validation of the multi-epitope vaccine. (A) Graphical illustration of the secondary structure prediction showing the presence of coil, helix and strand regions. (B) Tertiary structure after refinement showed in space-filling model. (C) Ramachandran plot of the 3D refined structure. (D) Z-score plot obtained from ProSA-web.

The tertiary structure was subjected to refinement using the GalaxyRefine server. The output showed five potential models. Model 1 (Fig. 3B) was classified as the best based on various parameters such as GDT-HA (0.9886), RMSD (0.292), MolProbity (2.364), clash score (24.4), and poor rotamers score (0.3). Therefore, this model was selected for further analysis. In this regard, the Ramachandran plot (Fig. 3C) showed that 86.4% of residues were located in most favoured regions, whereas the remaining residues were observed in additional allowed (11.7%), generously allowed (1.2%), and disallowed (0.6%) regions. In addition, the Z-score value (−2.35) (Fig. 3D) suggests that the vaccine structure is similar to native proteins of comparable size.

#### 3.4.3. Prediction of conformational BC epitopes

Three conformational BC epitopes (CE) were predicted using Ellipro (Fig. 4). These CE showed high probability scores—CE1: 0.914, CE2: 0.841 and CE3: 0.821, suggesting a considerable accessibility for antibodies (Fig. 4). Likewise, these results also confirm the immunogenic potential of the multi-epitope vaccine construct.

**Fig. 4.**
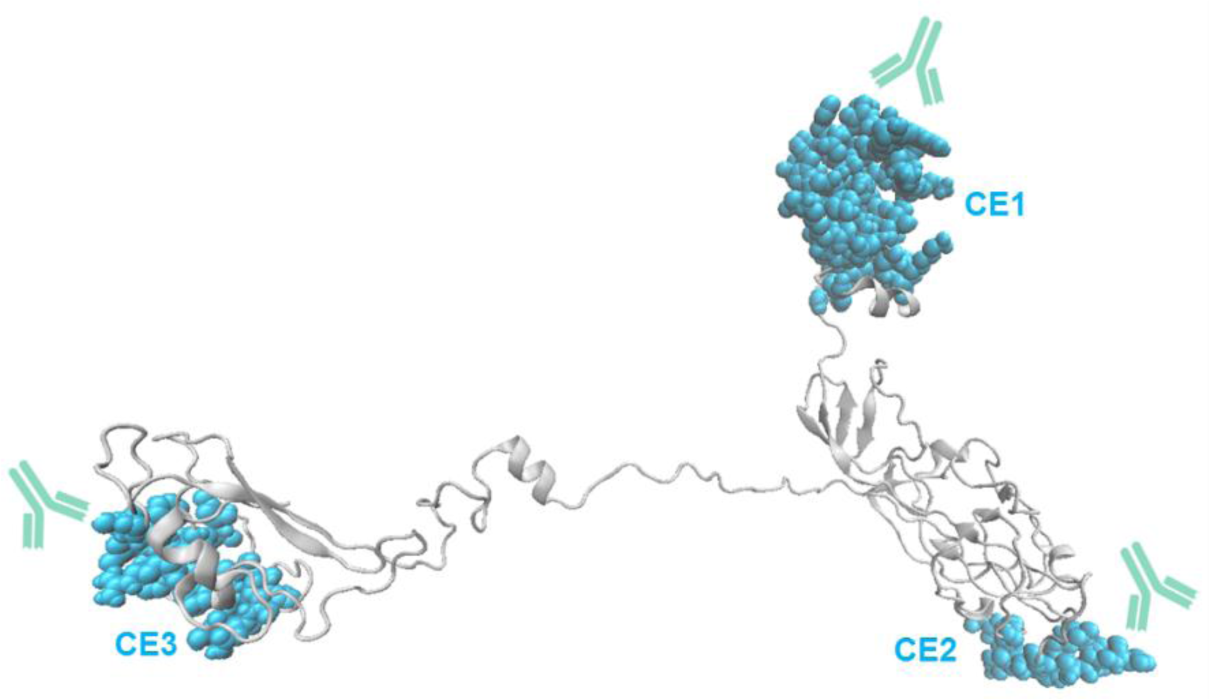
Best conformational B cell epitopes (CE) identified in the multi-epitope vaccine. These epitopes (cyan atoms) showed high probability scores (>0.8) suggesting a considerable accessibility for antibodies (showed in turquoise).

#### 3.4.4. TLR4-vaccine model interaction: Molecular docking

The output parameters of the TLR4*–*vaccine complex showed a suitable geometric shape complexity (29.237) and the interface area of the interaction (3271.40). Moreover, Fig. 5 clearly showed that the multi-epitope vaccine construct properly occupied the TLR4.

**Fig. 5.**
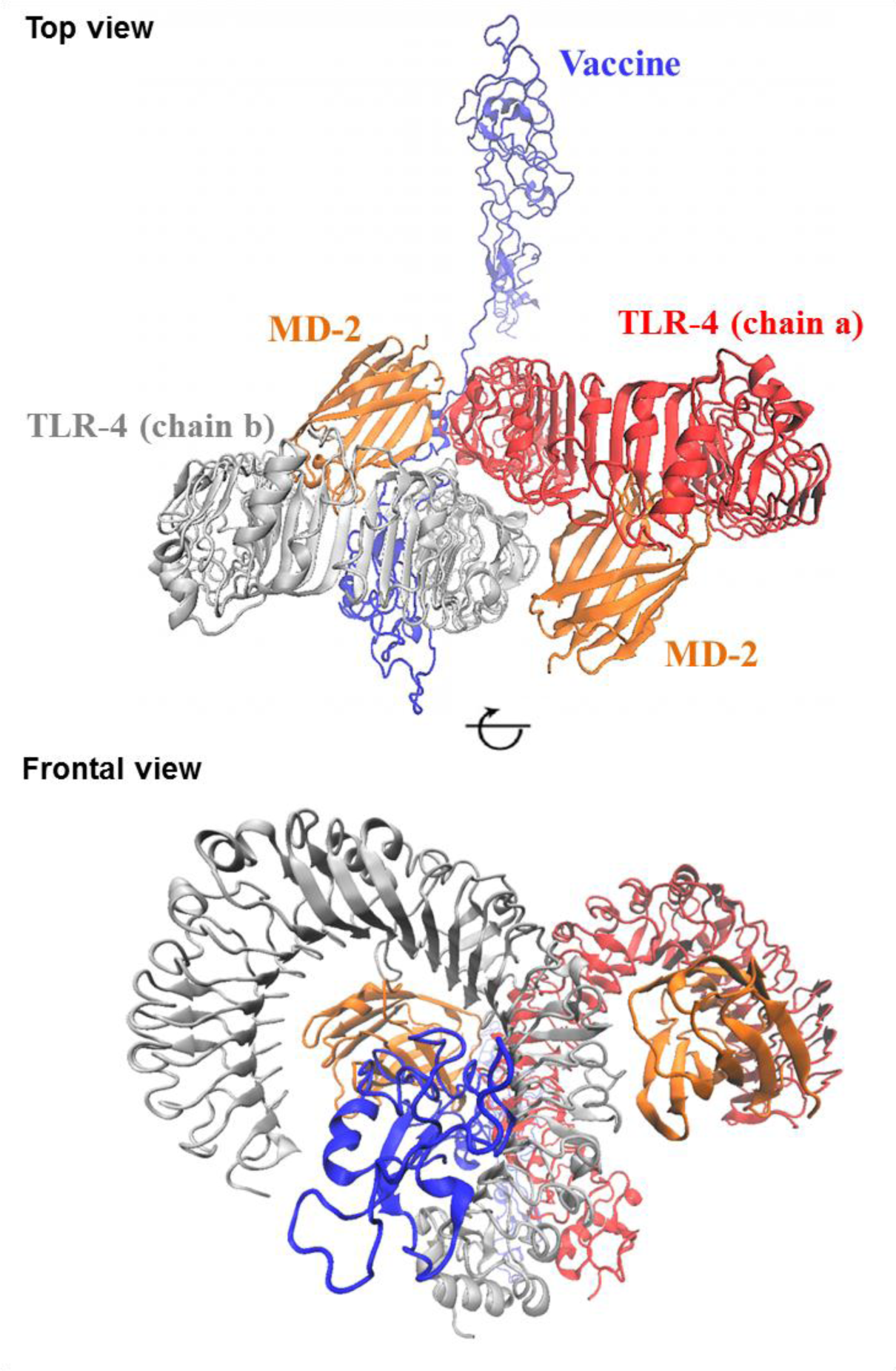
Docked complex of TLR4, MD-2 (myeloid differentiation factor 2), and the multi-epitope vaccine.

#### 3.4.5. Immune response simulations

The immune response simulations with the multi-epitope vaccine construct (3 doses given 4 weeks apart) showed cell-mediated and humoral responses. As expected, increased number and activity of Natural Killer (NK) cells—one of the first lines of defence against viruses [18], and macrophages were observed (Fig. 6). Regarding the adaptive immune response, CTC and THC populations showed a proliferative burst, effector cell generation, and a dramatic cell number contraction (Fig. 6). Importantly, IL-2, which is necessary for T cell activation and optimal proliferation [18], was amplified after each dose (Fig. 6). Moreover, the vaccine model increased BC and plasma cell populations, particularly immunoglobulin M (IgM) and IgG1 isotypes (Fig. 6). In this regard, titres of IgM, IgG1, and IgG2 were higher in the secondary and tertiary response compared to primary response (Fig. 6). Of note, immunogen concentrations decreased after antibody response (Fig. 6). Notably, repeated exposure with 12 injections (given 4 weeks apart) increased the IgG1 levels and stimulated CTC and THC populations (Fig. S1). Taken together, these results suggest that the multi-epitope vaccine could evoke and maximize both effector responses and immunological memory to SARS-CoV-2.

**Fig. 6.**
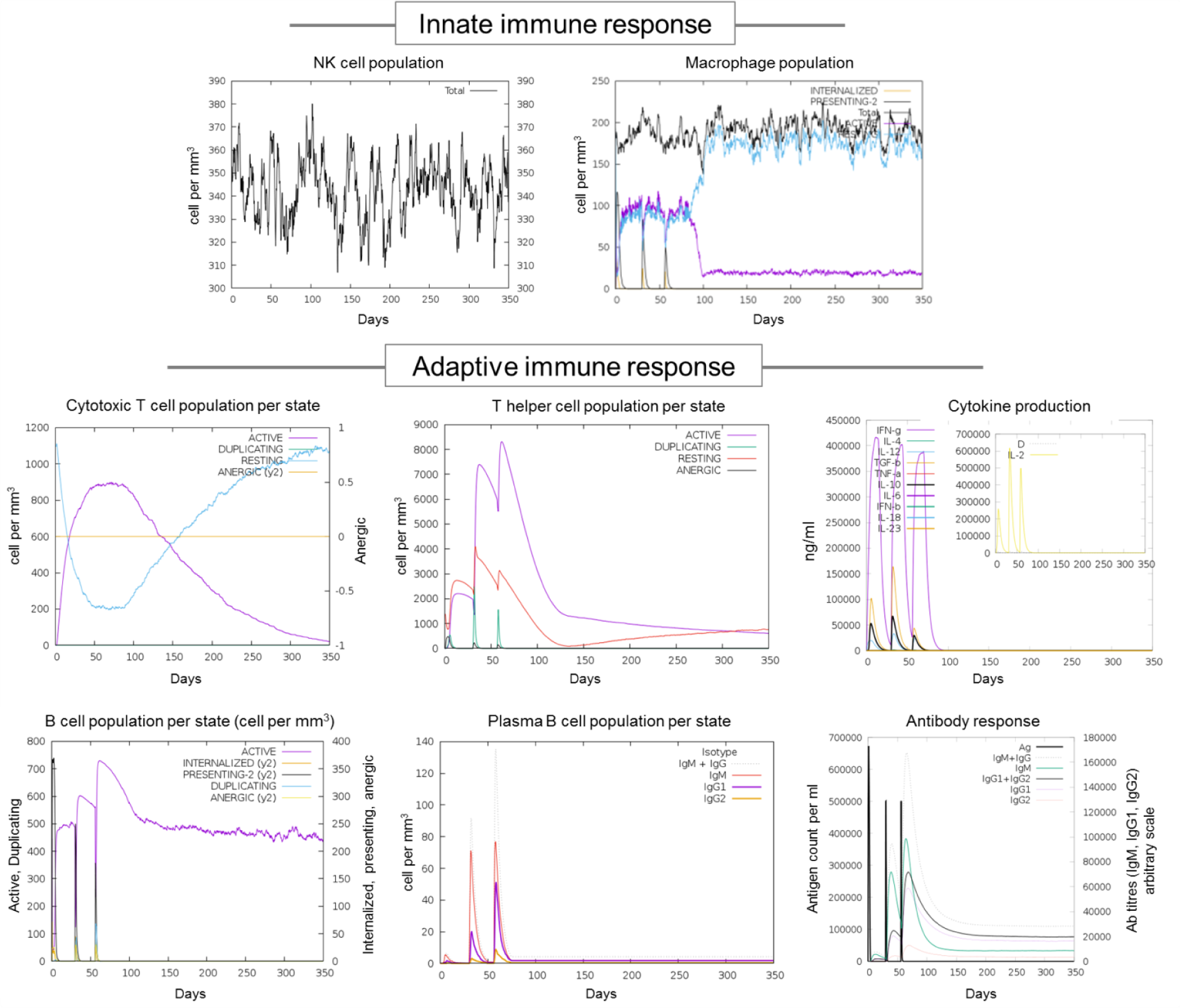
Immune response simulations with the multi-epitope vaccine (3 doses given 4 weeks apart).

#### 3.4.6. Population coverage

To investigate whether the multi-epitope vaccine may be used in different ethnic groups or globally, a population coverage analysis was performed. Remarkably, the multi-epitope vaccine construct showed high global population coverage: 99.69%. For instance, several countries with positive reports of COVID-19 (>6000 cases) [53], obtained the highest values, including, Australia, Brazil, Ecuador, Chile, China, France, Germany, India, Iran, Israel, Italy, Japan, Mexico, Morocco, Peru, Philippines, Russia, Singapore, South Korea, Spain, Sweden, USA, UK, etc., (Fig. 7).

**Fig. 7.**
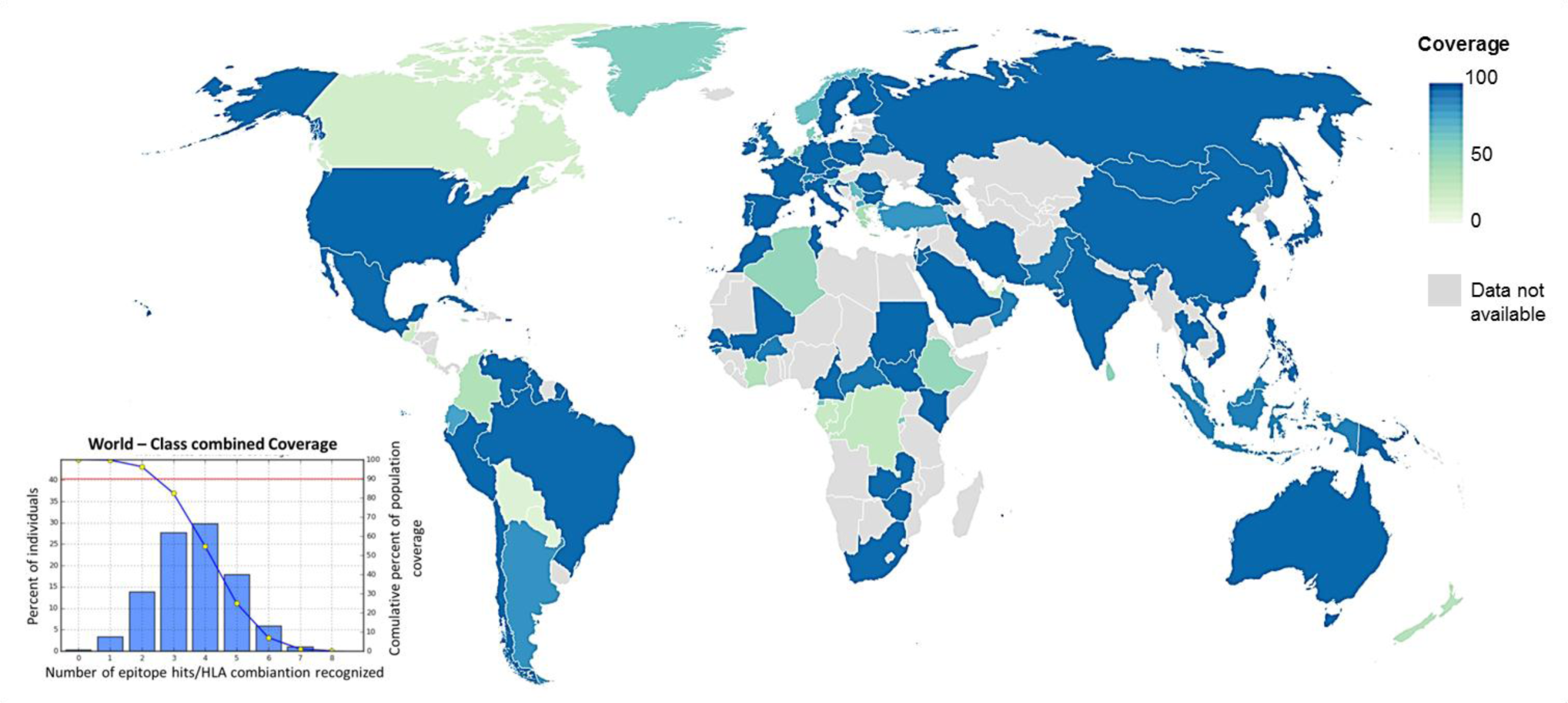
Combined HLA population coverage analysis of the multi-epitope vaccine against SARS-CoV-2.

## 4. Discussion

Immunoinformatics represents a valuable tool whereby the limitations in the selection of appropriate antigens and immunodominant epitopes may be overcome [17]. Previous *in silico*-based reports have shown that the SARS-CoV-2 S glycoprotein contains potential epitopes [8-11]. Therefore, researchers have recently attempted to design epitope-based vaccine candidates against SARS-CoV-2 [10, 11]. Despite these relevant contributions, one group only used T cell epitopes and did not include BC epitopes [10], which are fundamental players in antiviral immune response [18]. The second work, on the other hand, considered several viral membrane proteins, including the S glycoprotein, to identified probable T and BC epitopes [11]. Although the predicted epitopes showed good immunogenic potential, the vaccine does not target S glycoprotein RBM [11]. In the present study, highly potential B and T cell epitopes from the SARS-CoV-2 glycoprotein were predicted and the best selected to design a high-quality multi-epitope vaccine candidate. Remarkably, this vaccine harbours 2 epitopes (E19 in Table 1 and E10 in Table 2) that could evoke immune responses against SARS-CoV-2 RBM—the main responsible for virus entry into human cells [4,51], whereas 4 epitopes (E43, E44, E45, and E46 in Table 1) may direct the immune attack against other regions of SARS-CoV-2 RBD. These results are consistent with *in vitro* data that have demonstrated the antigenicity of the SARS-CoV-2 S glycoprotein [4].

The T cell epitopes included in the vaccine sequence accomplish with relevant requisites to design a suitable multi-epitope vaccine candidate. Firstly, they showed a marked antigenicity, immunogenicity, and lack of allergenic or toxic residues. Secondly, the THC epitopes were predicted as potent inductors of IFN-g—a crucial cytokine for CTC activation [18]. Thirdly, both CTC and THC epitopes properly interacted with the groove of HLA-I and HLA-II alleles, respectively, which is in agreement with other computer-based reports [54], thereby suggesting that the T cell epitopes identified and selected in the present study could be successfully presented in the context of HLA molecules.

The purpose of an adjuvant is to make a vaccine “detectable” for antigen-presenting cells such as dendritic cells [55]. In this regard, adjuvants approved or in clinical trials (NCT01609257) for virus-like particle-based vaccines are constituted by TLR4 agonists [55]. Here, the TLR4 adjuvant known as RS09 [33] was included in the multi-epitope vaccine sequence. The molecular docking simulation showed that the multi-epitope vaccine rightly interacts with this innate immune receptor in a similar way to previous works [35].

Notably, this study shows, by immunoinformatics simulations, the induction of both innate and adaptive responses to SARS-CoV-2. In this regard, NK cell and macrophage activation were detected, as well as high production of typical antibodies (IgM and IgG), cytokines (IFN-g and IL-2), and a proliferative burst of CTC and THC were observed after three injections. The generation and increase of plasma cells were also documented. Furthermore, B and T cell populations decreased along with immunogen levels. These data is comparable to previous investigations that have been focused on vaccine development against *Mycobacterium ulcerans* [46] and filarial diseases [56], as well as are in agreement, at least partially, with a recent study that demonstrated a positive correlation between robust CD4+ THC responses with anti-SARS-CoV-2 IgG and IgA titres of COVID-19 convalescent patients [57]. These immune responses were directed to the SARS-CoV-2 S glycoprotein [57].

This work was limited by A) the population coverage analysis did not include some countries, particularly from Africa, Central America, Eastern Europe, and Central Asia. This was mainly due to data not available concerning the HLA allele frequencies. Nevertheless, the highest population coverage was observed in several of the worst-hit countries by COVID-19 (e.g, Brazil, China, France, Italy, Iran, Peru, Spain, USA, etc.) [52]. B) This study did not explore whether the epitopes used for vaccine design are conserved in other beta-coronaviruses. However, former reports have already demonstrated that SARS-CoV-2 shares 79.5% and 50% sequence identity to SARS-CoV and MERS-CoV, respectively [58].

In summary, this study provides a novel multi-epitope vaccine built from high potential epitopes derived from the SARS-CoV-2 S glycoprotein. This immunoinformatics study suggests that such multi-epitope vaccine could activate and generate robust humoral and cell-mediated responses in a simultaneous manner against SARS-CoV-2, as well as the population coverage analysis indicates that it could be used globally. However, further rigorous *in vitro* and in *vivo* studies are imperative to confirm its immunogenic properties, safety, and efficacy, which—of course—would imply months, even years.

## Acknowledgments

I wish to thank the support of the Venezuelan Institute for Scientific Research (IVIC) – Venezuela, and Laboratory of Cellular and Molecular Pathology-IVIC.

## Founding

This research did not receive any specific grant from funding agencies in the public, commercial, or not-for-profit sectors.

---

**Fig. S1.**
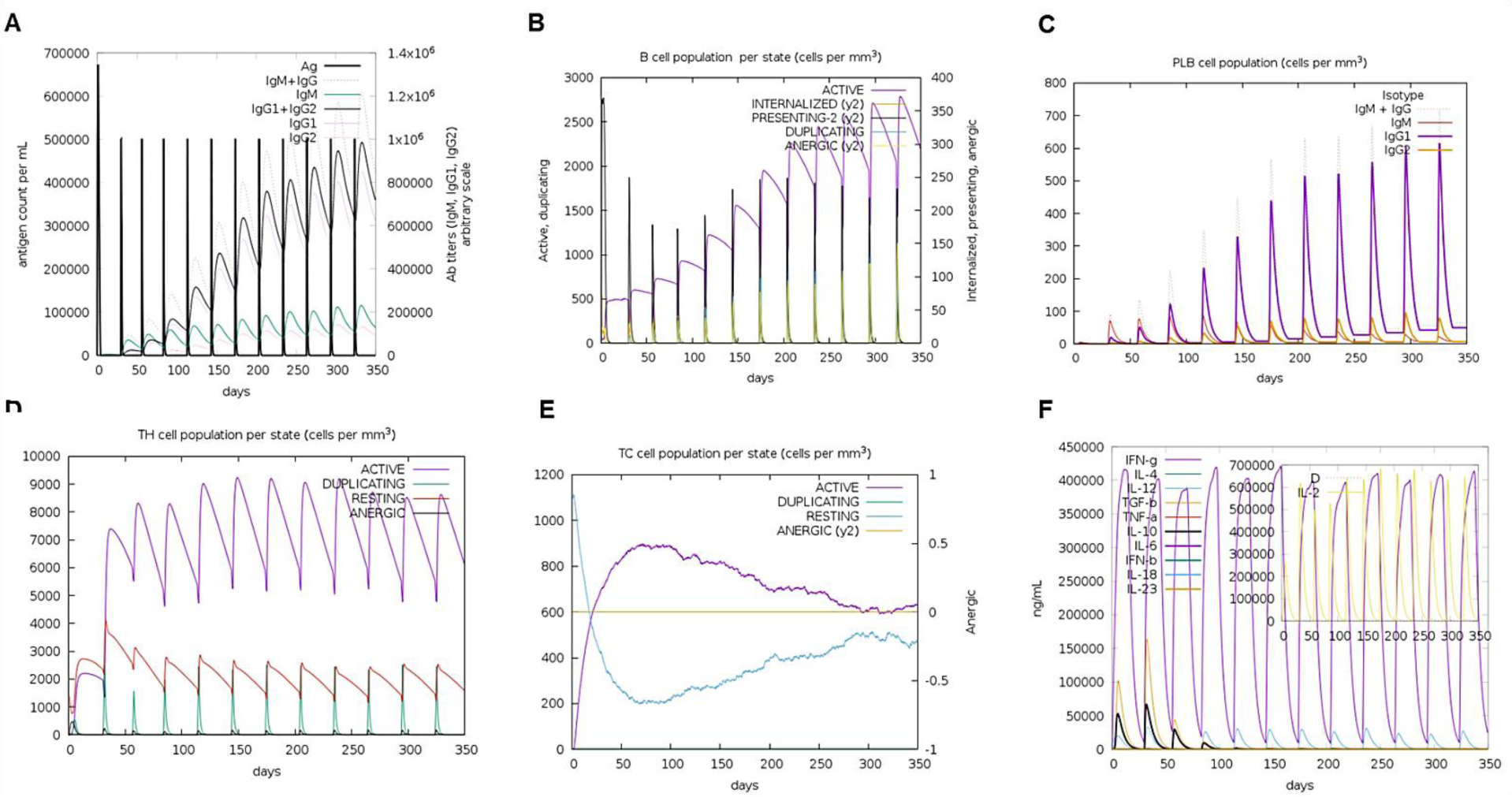
Immune response simulations with the multi-epitope vaccine (12 doses given 4 weeks apart). (A) B-cell activation and production of immunoglobulin. (B-C) Amount of B-cell and plasma cell populations. (D-E) T-helper cell and CTC cell populations per state after stimulation. (F) Cytokine levels induced after vaccine doses, including IL-2 (inset).

## References

[1] World Health Organization. Coronavirus disease (COVID-19) Pandemic. https://www.who.int/emergencies/diseases/novel-coronavirus-2019 (Accessed 19 May 2020).

[2] P. Zhou, X.L. Yang, X.G. Wang, B. Hu, L. Zhang, W. Zhang, et al., A pneumonia outbreak associated with a new coronavirus of probable bat origin, Nature. 579 (2020) 270–273, https://doi.org/10.1038/s41586-020-2012-7.

[3] Y. Jin, H. Yang, W. Ji, W. Wu, S. Chen, W. Zhang, G. Duan, Virology, Epidemiology, Pathogenesis, and Control of COVID-19, Viruses. 12 (2020) E372, https://doi.org/10.3390/v12040372.

[4] A.C. Walls, Y.J. Park, M.A. Tortorici, A. Wall, A.T. McGuire, D. Veesler, Structure, Function, and Antigenicity of the SARS-CoV-2 Spike Glycoprotein, Cell. 181 (2020) 281–292, https://doi.org/10.1016/j.cell.2020.02.058.

[5] F. Amanat, F. Krammer, SARS-CoV-2 Vaccines: Status Report, Immunity. 52 (2020) 583–589, https://doi.org/10.1016/j.immuni.2020.03.007.

[6] W.Y. Zhou, Y. Shi, C. Wu, W.J. Zhang, X.H. Mao, G. Guo, H.X. Li, Q.M. Zou, Therapeutic efficacy of a multi-epitope vaccine against Helicobacter pylori infection in BALB/c mice model, Vaccine. 27 (2009) 5013–5019, https://doi.org/10.1016/j.vaccine.2009.05.009.

[7] M.R. Dikhit, A. Kumar, S. Das, B. Dehury, A.K. Rout, F. Jamal, G.C. Sahoo, R.K. Topno, K. Pandey, V. Das, S. Bimal, P. Das, Identification of Potential MHC Class-II-Restricted Epitopes Derived from Leishmania donovani Antigens by Reverse Vaccinology and Evaluation of Their CD4+ T-Cell Responsiveness against Visceral Leishmaniasis, Front. Immunol. 8 (2017) 1763, https://doi.org/10.3389/fimmu.2017.01763.

[8] A. Grifoni, J. Sidney, Y. Zhang, R.H. Scheuermann, B. Peters, A. Sette, A Sequence Homology and Bioinformatic Approach Can Predict Candidate Targets for Immune Responses to SARS-CoV-2, Cell. Host. Microbe. 27(2020) 671–680.e2, https://doi.org/10.1016/j.chom.2020.03.002.

[9] G. Lucchese, Epitopes for a 2019-nCoV vaccine, Cell. Mol. Immunol. 17 (2020) 539–540, https://doi.org/10.1038/s41423-020-0377-z.

[10] M. Bhattacharya, A.R. Sharma, P. Patra, P. Ghosh, G. Sharma, B.C. Patra, S.S. Lee, C. Chakraborty, Development of epitope-based peptide vaccine against novel coronavirus 2019 (SARS-COV-2): Immunoinformatics approach, J. Med. Virol. (2020) 10.1002/jmv.25736. Advance online publication, https://doi.org/10.1002/jmv.25736.

[11] P. Kalita, A. Padhi, K. Zhang, T. Tripathi, Design of a peptide-based subunit vaccine against novel coronavirus SARS-CoV-2, Microb. Pathog. (2020) 104236 Advance online publication, https://doi.org/10.1016/j.micpath.2020.104236.

[12] World Health Organization. Biologicals: Nonclinical evaluation of vaccines. https://www.who.int/biologicals/vaccines/nonclinial_evaluation_of_vaccines/en/ (Accessed 19 May 2020).

[13] I. Dimitrov, L. Naneva, I. Doytchinova, I. Bangov, AllergenFP: allergenicity prediction by descriptor fingerprints, Bioinformatics. 30 (2014) 846–851, https://doi.org/10.1093/bioinformatics/btt619.

[14] S. Saha, G.P. Raghava, AlgPred: prediction of allergenic proteins and mapping of IgE epitopes, Nucleic Acids. Res. 34 (2006) W202–W209, https://doi.org/10.1093/nar/gkl343.

[15] S. Gupta, P. Kapoor, K. Chaudhary, A. Gautam, R. Kumar, Open Source Drug Discovery Consortium, G. P. Raghava, In silico approach for predicting toxicity of peptides and proteins, PloS one. 8 (2013) e73957, https://doi.org/10.1371/journal.pone.0073957.

[16] I.A. Doytchinova, D.R. Flower, VaxiJen: a server for prediction of protective antigens, tumour antigens and subunit vaccines, BMC bioinformatics. 8 (2007) 4, https://doi.org/10.1186/1471-2105-8-4.

[17] M. Sharma, F. Krammer, A. García-Sastre, S. Tripathi, Moving from Empirical to Rational Vaccine Design in the ‘Omics’ Era, Vaccines. 7 (2019) 89, https://doi.org/10.3390/vaccines7030089.

[18] J. Owen, J. Punt, S. Stranford, J. Pat, Kuby Immunology, Seventh ed., W.H. Freeman and Company, New York, 2013.

[19] Y. Kim, J. Ponomarenko, Z. Zhu, D. Tamang, P. Wang, J. Greenbaum, C. Lundegaard, A. Sette, O. Lund, P.E. Bourne, M. Nielsen, B. Peters, Immune epitope database analysis resource, Nucleic Acids. Res. 40 (2012) W525–W530, https://doi.org/10.1093/nar/gks438.

[20] S. Tenzer, B. Peters, S. Bulik, O. Schoor, C. Lemmel, M.M. Schatz, P.M. Kloetzel, H.G. Rammensee, H. Schild, H.G. Holzhütter, Modeling the MHC class I pathway by combining predictions of proteasomal cleavage, TAP transport and MHC class I binding, Cell. Mol. Life Sci. 62 (2005) 1025–1037, https://doi.org/10.1007/s00018-005-4528-2.

[21] M. Moutaftsi, B. Peters, V. Pasquetto, D.C. Tscharke, J. Sidney, H.H. Bui, H. Grey, A. Sette, A consensus epitope prediction approach identifies the breadth of murine T(CD8+)-cell responses to vaccinia virus, Nat. Biotechnol. 24 (2006) 817–819, https://doi.org/10.1038/nbt1215.

[22] I. Hoof, B. Peters, J. Sidney, L.E. Pedersen, A. Sette, O. Lund, S. Buus, M. Nielsen, NetMHCpan, a method for MHC class I binding prediction beyond humans. Immunogenetics, 61(2009) 1–13, https://doi.org/10.1007/s00251-008-0341-z.

[23] P. Wang, J. Sidney, C. Dow, B. Mothé, A. Sette, B. Peters, A systematic assessment of MHC class II peptide binding predictions and evaluation of a consensus approach, PLoS Comput. Biol. 4 (2008) e1000048, https://doi.org/10.1371/journal.pcbi.1000048.

[24] M. Nielsen, C. Lundegaard, T. Blicher, B. Peters, A. Sette, S. Justesen, S. Buus, O. Lund,. Quantitative predictions of peptide binding to any HLA-DR molecule of known sequence: NetMHCIIpan, PLoS Comput. Biol. 4 (2008) e1000107, https://doi.org/10.1371/journal.pcbi.1000107.

[25] M. Nielsen, C. Lundegaard, O. Lund, Prediction of MHC class II binding affinity using SMM-align, a novel stabilization matrix alignment method, BMC bioinformatics. 8 (2007) 238, https://doi.org/10.1186/1471-2105-8-238.

[26] S.K. Dhanda, P. Vir, G.P. Raghava, Designing of interferon-gamma inducing MHC class-II binders, Biol. Direct. 8 (2013) 30, https://doi.org/10.1186/1745-6150-8-30.

[27] S. Saha, G.P. Raghava, Prediction of continuous B-cell epitopes in an antigen using recurrent neural network, Proteins. 65 (2006) 40–48, https://doi.org/10.1002/prot.21078.

[28] B. Manavalan, R.G. Govindaraj, T.H. Shin, M.O. Kim, G. Lee, iBCE-EL: A New Ensemble Learning Framework for Improved Linear B-Cell Epitope Prediction, Front. Immunol. 9 (2018) 1695, https://doi.org/10.3389/fimmu.2018.01695.

[29] M.C. Jespersen, B. Peters, M. Nielsen, P. Marcatili, BepiPred-2.0: improving sequence-based B-cell epitope prediction using conformational epitopes, Nucleic Acids. Res. 45 (2017) W24–W29, https://doi.org/10.1093/nar/gkx346.s

[30] J. Maupetit, P. Derreumaux, P. Tuffery, A fast and accurate method for large-scale de novo peptide structure prediction, J. Comput. Chem. 31 (2010) 726–38, https://doi.org/10.1002/jcc.21365.

[31] D. Schneidman-Duhovny, Y. Inbar, R. Nussinov, H.J. Wolfson, PatchDock and SymmDock: servers for rigid and symmetric docking, Nucleic Acids. Res. 33(2005) W363–W367, https://doi.org/10.1093/nar/gki481.

[32] W. Humphrey, A. Dalke, K. Schulten, VMD –Visual Molecular Dynamics, J. Molec. Graph. 14 (1996) 33–38, https://doi.org/10.1016/0263-7855(96)00018-5.

[33] A. Shanmugam, S. Rajoria, A.L. George, A. Mittelman, R. Suriano, R.K. Tiwari, Synthetic Toll like receptor-4 (TLR-4) agonist peptides as a novel class of adjuvants, PloS one. 7 (2012) e30839, https://doi.org/10.1371/journal.pone.0030839.

[34] S.N. Lester, K. Li, Toll-like receptors in antiviral innate immunity, J. Mol. Biol. 426 (2014) 1246–1264, https://doi.org/10.1016/j.jmb.2013.11.024.

[35] R.K. Pandey, T.K. Bhatt, V.K. Prajapati, Novel Immunoinformatics Approaches to Design Multi-epitope Subunit Vaccine for Malaria by Investigating Anopheles Salivary Protein, Sci. Rep. 8 (2018) 1125, https://doi.org/10.1038/s41598-018-19456-1.

[36] M.R. Wilkins, E. Gasteiger, A. Bairoch, J.C. Sanchez, K.L. Williams, R.D. Appel, D.F. Hochstrasser, Protein identification and analysis tools in the ExPASy server, Methods Mol Biol. 112 (1999) 531–552, https://doi.org/10.1385/1-59259-584-7:531.

[37] C.N. Magnan, A. Randall, P. Baldi, SOLpro: accurate sequence-based prediction of protein solubility, Bioinformatics. 25 (2009) 2200–2207, https://doi.org/10.1093/bioinformatics/btp386.

[38] D.T. Jones, Protein secondary structure prediction based on position-specific scoring matrices, J. Mol. Biol. 292 (1999) 195–202, https://doi.org/10.1006/jmbi.1999.3091.

[39] D. Buchan, D.T. Jones, The PSIPRED Protein Analysis Workbench: 20 years on, Nucleic Acids. Res. 47 (2019) W402–W407, https://doi.org/10.1093/nar/gkz297.

[40] W.H. Shin, G. R. Lee, L. Heo, H. Lee, C. Seok, Prediction of Protein Structure and Interaction by GALAXY protein modeling programs, Bio. Design. 2 (2014) 1–11.

[41] J. Ko, H. Park, L. Heo, C. Seok, GalaxyWEB server for protein structure prediction and refinement, Nucleic Acids. Res. 40 (2012) W294–W297, https://doi.org/10.1093/nar/gks493.

[42] R.A. Laskowski, M.W. MacArthur, D.S. Moss, J.M. Thornton, PROCHECK - a program to check the stereochemical quality of protein structures, J. App. Cryst. 26 (1993) 283–291.

[43] M. Wiederstein, M.J. Sippl, ProSA-web: interactive web service for the recognition of errors in three-dimensional structures of proteins, Nucleic acids research. 35(2007), W407–W410, https://doi.org/10.1093/nar/gkm290.

[44] J. Ponomarenko, H.H. Bui, W. Li, N. Fusseder, P.E. Bourne, A. Sette, B. Peters, ElliPro: a new structure-based tool for the prediction of antibody epitopes, BMC bioinformatics. 9 (2008) 514, https://doi.org/10.1186/1471-2105-9-514.

[45] N. Rapin, O. Lund, M. Bernaschi, F. Castiglione, Computational immunology meets bioinformatics: the use of prediction tools for molecular binding in the simulation of the immune system, PloS one. 5(2010) e9862, https://doi.org/10.1371/journal.pone.0009862.

[46] Z. Nain, M.M. Karim, M.K. Sen, U.K. Adhikari, Structural basis and designing of peptide vaccine using PE-PGRS family protein of Mycobacterium ulcerans-An integrated vaccinomics approach, Mol. Immunol. 120 (2020) 146–163, https://doi.org/10.1016/j.molimm.2020.02.009.

[47] H.H. Bui, J. Sidney, K. Dinh, S. Southwood, M.J. Newman, A. Sette, Predicting population coverage of T-cell epitope-based diagnostics and vaccines, BMC bioinformatics. 7 (2006) 153, https://doi.org/10.1186/1471-2105-7-153.

[48] RStudio Team (2015). RStudio: Integrated Development for R. RStudio, Inc., Boston, MA URL http://www.rstudio.com/.

[49] J.T. Ortega, M.L. Serrano, F.H. Pujol, H.R. Rangel, Role of changes in SARS-CoV-2 spike protein in the interaction with the human ACE2 receptor: An in silico analysis, EXCLI. J. 19 (2020) 410–417, https://doi.org/10.17179/excli2020-1167.

[50] J. Lan, J. Ge, J. Yu, S. Shan, H. Zhou, S. Fan, Q. Zhang, X. Shi, Q. Wang, L. Zhang, X. Wang, Structure of the SARS-CoV-2 spike receptor-binding domain bound to the ACE2 receptor, Nature. 581(7807) (2020) 215–220, https://doi.org/10.1038/s41586-020-2180-5.

[51] J. Shang, G. Ye, K. Shi, Y. Wan, C. Luo, H. Aihara, Q. Geng, A. Auerbach, F. Li, Structural basis of receptor recognition by SARS-CoV-2, Nature. (2020) 10.1038/s41586-020-2179-y. https://doi.org/10.1038/s41586-020-2179-y.

[52] D. Duhovny, R. Nussinov, H.J. Wolfson, Efficient Unbound Docking of Rigid Molecules, in Gusfield et al. (Eds.), Proceedings of the 2’nd Workshop on Algorithms in Bioinformatics(WABI) Rome, Italy, Lecture Notes in Computer Science, Springer Verlag, 2452, pp. 185–200.

[53] World Health Organization. Coronavirus disease (COVID-2019) situation reports. https://www.who.int/emergencies/diseases/novel-coronavirus-2019/situation-reports/ (Accessed 19 May 2020).

[54] V. Baruah, S. Bose, Immunoinformatics-aided identification of T cell and B cell epitopes in the surface glycoprotein of 2019-nCoV, J. Med. Virol. 92 (2020) 495–500, https://doi.org/10.1002/jmv.25698.

[55] V. Cimica, J.M. Galarza, Adjuvant formulations for virus-like particle (VLP) based vaccines, Clin. Immunol. 183 (2017) 99–108, https://doi.org/10.1016/j.clim.2017.08.004.

[56] R.A. Shey, S.M. Ghogomu, K.K. Esoh, N.D. Nebangwa, C.M. Shintouo, N.F. Nongley, B.F. Asa, F.N. Ngale, L. Vanhamme, J. Souopgui, In-silico design of a multi-epitope vaccine candidate against onchocerciasis and related filarial diseases, Sci. Rep. 9 (2019) 4409, https://doi.org/10.1038/s41598-019-40833-x.

[57] A. Grifoni, D. Weiskopf, S.I. Ramirez, J. Mateus, J.M. Dan, C.R. Moderbacher, S.A. Rawlings, A. Sutherland, L. Premkumar, R.S. Jadi, D. Marrama, A.M. de Silva, A. Frazier, A. Carlin, J.A. Greenbaum, B. Peters, F. Krammer, D.M. Smith, S. Crotty, A. Sette, Targets of T cellresponses to SARS-CoV-2 coronavirus in humans with COVID-19 disease and unexposed individuals, Cell. (2020), doi: https://doi.org/10.1016/j.cell.2020.05.015.

[58] R. Lu, X. Zhao, J. Li, P. Niu, B. Yang, H. Wu, W. Wang, H. Song, B. Huang, N. Zhu, Y. Bi, X. Ma, F. Zhan, L. Wang, T. Hu, H. Zhou, Z. Hu, W. Zhou, L. Zhao, J. Chen, Y. Meng, J. Wang, Y. Lin, J. Yuan, Z. Xie, J. Ma, W.J. Liu, D. Wang, W. Xu, E.C. Holmes, G.F. Gao, G. Wu, W. Chen, W. Shi, W. Tan,. Genomic characterisation and epidemiology of 2019 novel coronavirus: implications for virus origins and receptor binding, Lancet. 395 (10224) (2020) 565–574, https://doi.org/10.1016/S0140-6736(20)30251-8.

